# Overcome Prostate Cancer Resistance to Immune Checkpoint Therapy with Ketogenic Diet-Induced Epigenetic Reprogramming

**DOI:** 10.1101/2023.08.07.552383

**Authors:** Sean Murphy, Sharif Rahmy, Dailin Gan, Yini Zhu, Maxim Manyak, Jun Li, Xuemin Lu, Xin Lu

## Abstract

Advanced prostate cancer (PCa) is overwhelmingly resistant to immune checkpoint blockade (ICB) therapy, representing a formidable clinical challenge. In this study, we developed a syngeneic murine PCa model with acquired ICB resistance. Using this model, synergistic efficacy was achieved by combining anti-PD1 and anti-CTLA4 antibodies with histone deacetylase inhibitor (HDACi) vorinostat, a cyclic ketogenic diet (CKD), or supplementation of ketone body β-hydroxybutyrate (BHB, endogenous HDACi) via 1,3-butanediol-admixed food. CKD and BHB supplementation delayed PCa tumors as monotherapy, and both BHB and adaptive immunity are required for the anti-tumor activity of CKD. Single-cell transcriptomic and proteomic profiling revealed that the HDACi and ketogenesis-enhanced ICB therapy involves cancer-cell-intrinsic (upregulated MHC class I molecules) and extrinsic mechanisms (CD8^+^ T cell chemoattraction, M1/M2 macrophage rebalancing, monocyte differentiation toward antigen presenting cells, and diminished neutrophils). Overall, these findings underscore the potential of using HDACi and optimized KD to enhance ICB therapy for PCa.

## Introduction

Prostate cancer (PCa) is the second most commonly diagnosed malignancy and the fifth leading cause of cancer mortality for men worldwide ^1^. A handful of targeted therapy drugs have been approved to treat advanced PCa, but only offer modest survival benefits ^1^. Therefore, new therapeutics are highly desired. Immune checkpoint blockade (ICB) using antibodies against cytotoxic-T-lymphocyte-associated protein 4 (CTLA4) or programmed cell death 1/ programmed cell death 1 ligand 1 (PD1/PD-L1) generates durable therapeutic responses in a significant subset of patients across various cancer types ^2^. However, advanced PCa shows overwhelming *de novo* resistance to ICB ^3^. PCa typically features a moderate mutation burden and a suppressed tumor immune microenvironment (TIME), posing unique challenges for sensitizing PCa to ICB therapy.

One promising immunotherapy sensitization approach is epigenetic drugs targeting histone deacetylation ^4^. Histone acetylation is controlled by the balance between histone acetyltransferases and histone deacetylases (HDACs). By inhibiting HDAC activity and modulating gene expression, HDAC inhibitors (HDACi) remain promising anti-cancer drugs, including targeting PCa ^5^. Besides the direct effect on cancer cells, HDACi have diverse immune-modulatory activities. An intact immune system is required for the robust anti-cancer effects of class I/II HDACi vorinostat and panobinostat in murine cancer models ^6^. The immune-modulatory activity of HDACi encompasses both cancer-cell-intrinsic and extrinsic mechanisms. On the one hand, both class I-specific and pan-HDACi increase the cancer cell immunogenicity through upregulation of tumor-associated antigens and the components of the antigen processing machinery including TAP-1, TAP-2, and MHC class I and class II molecules (MHC-I, MHC-II) ^7, 8^. *In vitro* experiments show that HDACi-treated PCa cells are more prone to killing by CD8^+^ T cells ^9^; however, *in vivo* evidence is lacking. On the other hand, HDACi has profound impact on immune populations in the TIME, including dampening immunosuppressive cells such as regulatory T cells (Treg) and myeloid-derived suppressor cells (MDSCs) ^10, 11^, and biasing the T cell stimulatory function of dendritic cells (DCs) toward Th1 or Th2 polarization ^12, 13^. Translation of these findings to early phase clinical trials of combining entinostat and high-dose interleukin 2 (IL2) in treating renal cell carcinoma showed promising clinical activity associated with decreased Treg ^14^. Vorinostat is an HDACi approved by FDA to treat cutaneous T cell lymphoma ^15^. Enhanced anti-tumor effect by combining vorinostat and various types of immunotherapy was reported in mammary, renal and colorectal tumor models ^16^. Whether vorinostat effectively sensitizes syngeneic PCa models to ICB therapy remains to be determined.

Dietary interventions, including the ketogenic diet (KD), are emerging metabolic approaches to enhancing cancer therapy ^17^. KD consists of high fat, moderate to low protein, and very low carbohydrates, forcing the body to burn fat rather than glucose for energy ^18^. Clinical use of KD includes treating childhood epilepsy ^19^. KD was recently found to extend the longevity of mice ^20, 21^. KD is not approved to treat cancer, but it was postulated that KD could be a potential dietary manipulation for exploiting inherent oxidative metabolic differences between cancer cells and normal cells to improve standard therapeutic outcomes ^18^. The effect of KD in animal tumor models are varied with an overall moderate anti-tumor activity ^22^. Fatty acid oxidation of ingested KD in the liver produces ketone bodies, with β-hydroxybutyrate (BHB) being the most abundant one ^23^. Once taken up by a target tissue, BHB is converted back to acetyl-CoA to feed the tricarboxylic acid (TCA) cycle for oxidation and ATP production. In addition to being a carrier of energy, BHB functions as a signaling metabolite and affects gene expression and cellular functions through several mechanisms ^23^. Interestingly, BHB is an endogenous pan-HDACi and modulates gene expression and function by inhibiting HDAC1 & 3 (class I) and HDAC4 (class II) ^24^. The immune-modulatory activity of BHB started to be appreciated recently ^25^.

In this study, we generated an isogeneic murine PCa cell series containing sensitive and resistant lines to anti-PD1 and anti-CTLA4 ICB. Using this model, we demonstrated synergistic efficacy by combining ICB with HDACi vorinostat, a cyclic ketogenic diet (CKD), or BHB supplementation in eradicating ICB-resistant tumors. Through functional experiments and single-cell technologies mass cytometry (CyTOF) and single-cell RNA-sequencing (scRNA-seq), we found that both cancer-cell-intrinsic (augmented MHC-I and antigen presentation) and extrinsic mechanisms (CD8^+^ T cell chemoattraction, M1/M2 macrophage rebalancing, influx and differentiation of monocytes toward antigen presenting cells, and diminished neutrophils) underlie the combinatorial efficacy.

## Results

### Development of ICB-sensitive and ICB-resistant syngeneic PCa cell lines

Previously, we developed murine PCa cell lines from *PB-Cre^+^ Pten^L/L^ p53^L/L^ Smad4^L/L^* genetically engineered mouse model of metastatic PCa ^26^. These PPS cell lines grew in C57BL/6 mice after subcutaneous inoculation. We focused on one line, PPS-6239, and tested the effect of anti-PD1 and anti-CTLA4 treatments as single agents or in combination on established tumors (∼100mm^3^). PPS-6239 tumors showed partial response to anti-PD1 or anti-CTLA4 alone (10% cure rate in each case) but substantial response to the combination (80% cure rate, **Fig. 1A**). The mice with tumors either resected surgically or eradicated through treatment in each cohort were subsequently challenged with PPS-6239 or an irrelevant syngeneic PCa cell line RM9 (Ras/Myc-transformed). RM9 grew in all mice tested, but PPS-6239 showed a distinct pattern dependent on the treatment history: mice treated with isotype IgG displayed no rejection, mice treated with anti-PD1 or anti-CTLA4 displayed moderate rejection, and mice treated with combination therapy rejected all inoculation attempts (**Fig. 1B**). This result demonstrates that effective immunotherapy targeting PPS-6239 generated enduring and specific memory immunity.

**Figure 1.**
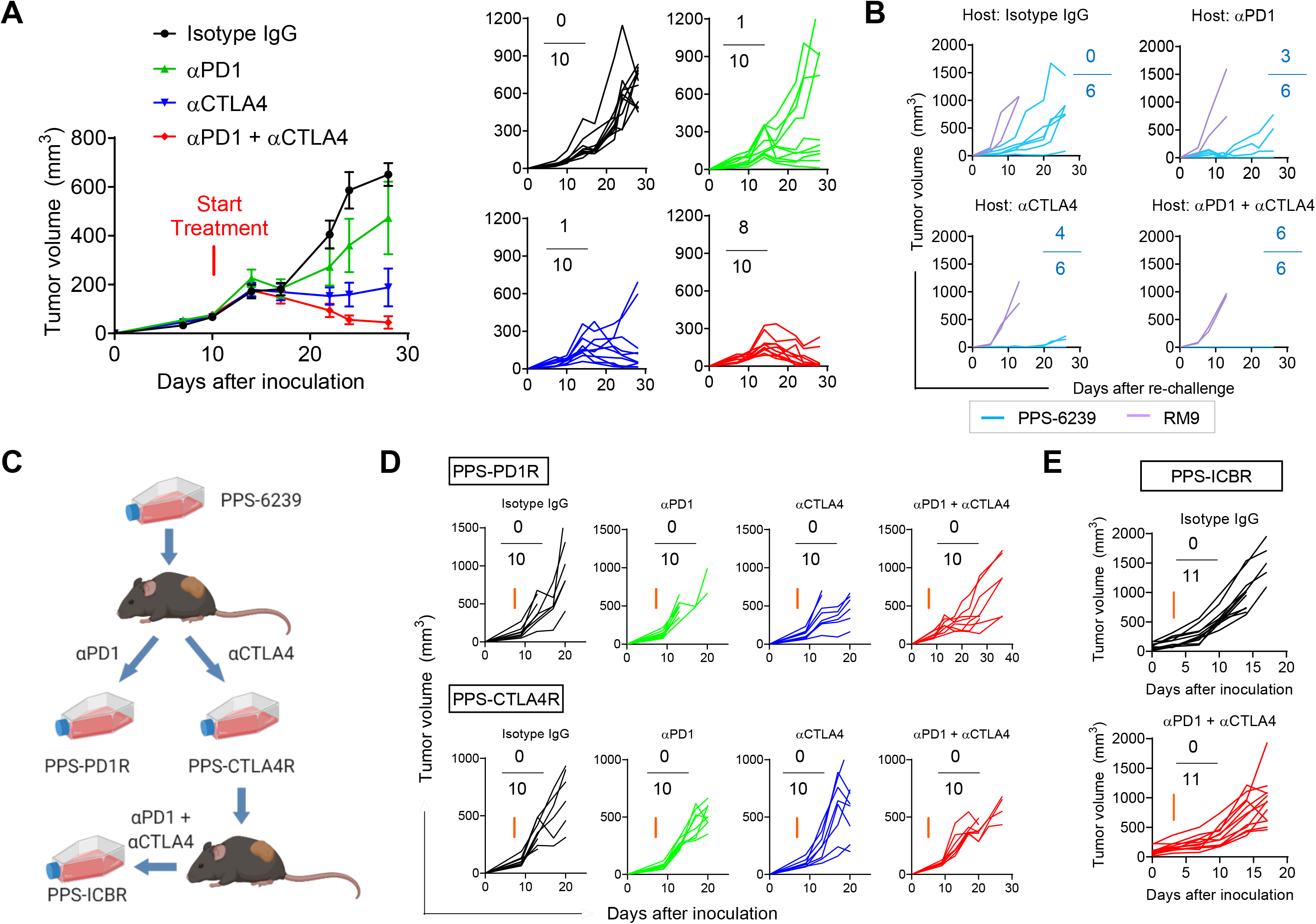
Development of ICB-sensitive and ICB-resistant syngeneic PCa cell lines. (A) Response of PPS-6239 tumors to single or combination ICB therapy, with both averaged and individual tumor volumes plotted. N=10/group. (B) Individual growth curves of PPS-6239 tumors or RM9 tumors in C57BL/6 hosts previously tumor-bearing with PPS-6239 and treated with the indicated antibodies. N=6/group. (C) The relationship of PPS-6239 and *in vivo* derivatives PPS-PD1R, PPS-CTLA4R and PPS-ICBR. (D) Response of PPS-PD1R and PPS-CTLA4R tumors to single or combination ICB therapy, with individual tumor volumes plotted. N=10/group. (E) Response of PPS-ICBR tumors to isotype IgG or combined αPD1+αCTLA4 antibodies, with individual tumor volumes plotted. N=11/group. In (A), (B), (D) and (E), red arrows denote the treatment start timepoint, and ratios represent the cured tumors over total tumors (i.e., complete response rate). In (A), data represent mean ± SEM.

In order to generate ICB-resistant variants of PPS-6239, we digested the rare tumors resistant to anti-PD1 or anti-CTLA4 single therapy and developed cell lines PPS-PD1R and PPS-CTLA4R, respectively (**Fig. 1C**). Both lines gained resistance to anti-PD1, anti-CTLA4 and the combination (0% cure rate in all groups, **Fig. 1D**). From one of the PPS-CTLA4R-bearing mice treated with the combination therapy, we further developed a subline PPS-ICBR (**Fig. 1C**), which exhibited resistance to anti-PD1 plus anti-CTLA4 therapy (0% cure rate, **Fig. 1E**). Consistent with the opposite response of PPS-6239 and PPS-ICBR to the dual ICB therapy, PPS-6239 tumors had massive infiltration of CD8^+^ T cells under the therapy. In contrast, despite the therapy, PPS-ICBR tumors were devoid of CD8+ T cells. The PPS-6239, PPS-PD1R, PPS-CTLA4R and PPS-ICBR series of isogenic cell lines provide a valuable model to study the mechanism of ICB resistance and the strategy to overcome it.

### Vorinostat sensitized PPS-ICBR tumors to ICB therapy

Consistent with the critical role of antigen presentation by MHC-I in tumor immunity, H2-K^b^ and H2-D^b^ were significantly downregulated in the resistant sublines (especially PPS-ICBR) compared with PPS-6239 (**Fig. 2A**). To assess whether HDACi could revert gene expression associated with ICB resistance, we profiled the transcriptome of PPS-ICBR treated with DMSO or 5µM vorinostat for 12 hours *in vitro*. Among the differentially expressed genes, MHC-encoded genes were significantly upregulated by vorinostat, including *H2-K1* and *H2-D1* (**Fig. 2B**). H2-K^b^ protein level was restored by 5μM vorinostat (**Fig. 2C**). Chromatin immunoprecipitation followed by quantitative PCR (ChIP-qPCR) using Histone H3 acetyl Lys9 (H3K9Ac) antibody showed augmented histone acetylation of the *H2-K1* locus in vorinostat-treated cells compared with DMSO-treated cells (**Fig. 2D**). KEGG pathway enrichment analysis showed that vorinostat-upregulated genes (FDR<0.05, Vo/DMSO fold change>3) enriched for immune-related pathways, such as TNF signaling, IL-17 signaling, and cytokine-cytokine receptor interaction (**Fig. 2E**), consistent with the immune-modulatory activities of HDACi. By contrast, vorinostat-downregulated genes (FDR<0.05, DMSO/Vo fold change>3) enriched for proliferation-related pathways such as DNA replication and cell cycle (**Fig. 2E**), consistent with the cytostatic activity of HDACi. When PPS-ICBR cells were transduced with lentivirus encoding shRNA targeting *Hdac1*, transient *Hdac1* knockdown upregulated H-2K^b^ and H-2D^b^ (**Fig. 2F**), but stable selection with puromycin resulted in no viable cells, suggesting that PPS-ICBR cells could not tolerate continuous Hdac1 loss.

**Figure 2.**
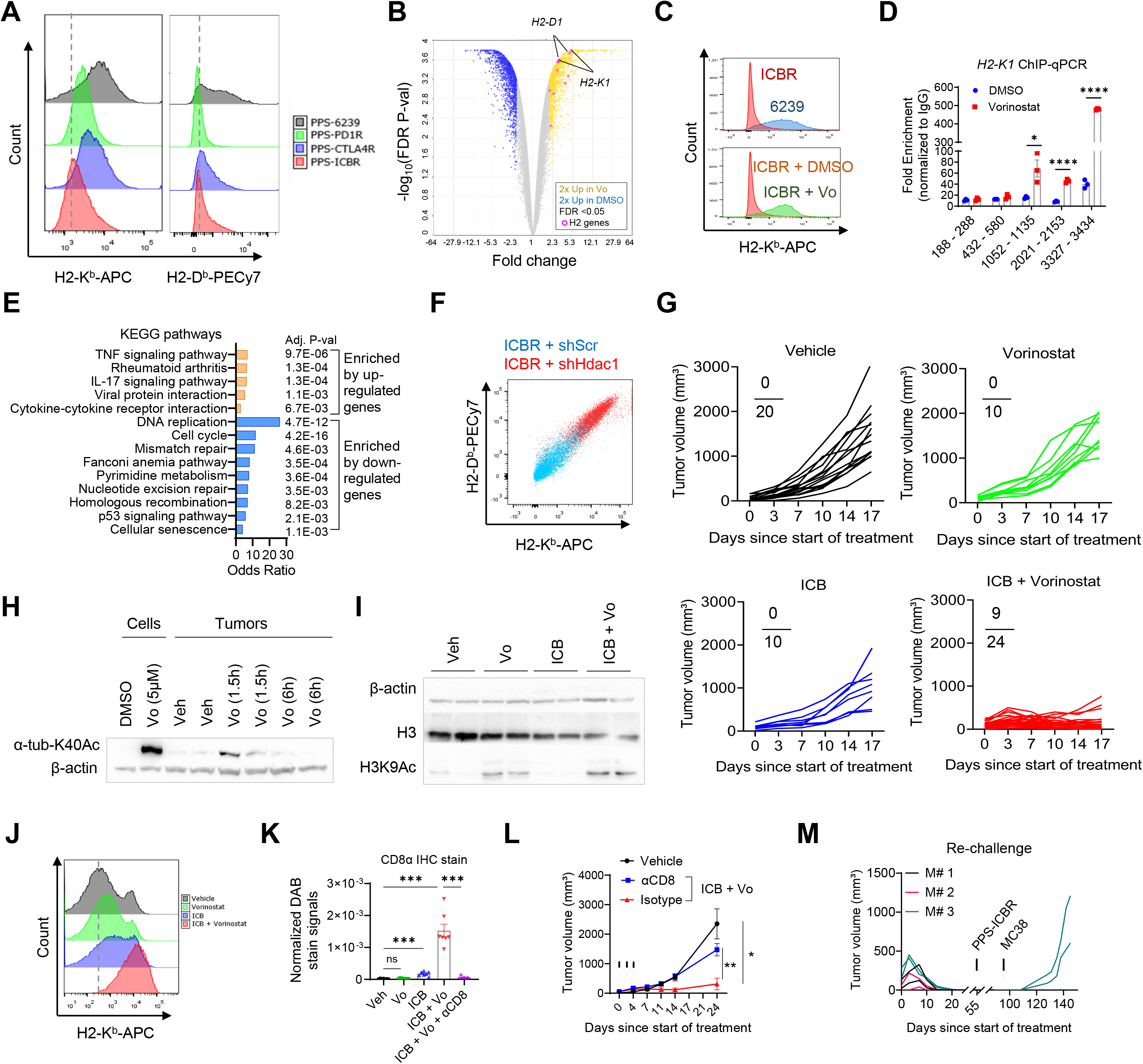
Vorinostat sensitized PPS-ICBR tumors to ICB therapy. (A) Flow cytometry histograms of H2-K^b^ and H2-D^b^ for the four cell lines. (B) Volcano plot for differentially expressed genes in PPS-ICBR cells treated with 5 µM vorinostat or DMSO for 12h. Cutoff thresholds and two MHC-I genes were indicated. (C) Flow cytometry histograms of H2-K^b^ for PPS-ICBR cells treated with 5 µM vorinostat or DMSO for 12h. (D) H3K9Ac ChIP-qPCR of the *H2-K1* locus in PPS-ICBR cells treated with 5µM vorinostat or DMSO. N=3/condition. (E) Top enriched KEGG pathways for genes upregulated or downregulated by vorinostat in PPS-ICBR cells. (F) Flow cytometry plot of H2-K^b^ and H2-D^b^ for PPS-ICBR cells transduced with control shRNA (scScr) or *Hdac1* shRNA (shHdac1). (G) Individual growth curves of PPS-ICBR tumors in mice treated with vehicle (n=20), vorinostat (n=10), ICB (anti-PD1+anti-CTLA4, n=10), or ICB+vorinostat (n=24). Complete response ratios were labeled. (H) Western blot of α-tub-K40Ac and β-actin of PPS-ICBR cells or tumors treated with vehicle or vorinostat. (I) Western blot of H3K9Ac, total H3 and β-actin of PPS-ICBR tumors treated with vehicle, vorinostat, ICB, or ICB+vorinostat. (J) Flow cytometry histograms of H2-K^b^ for PCa cells in the dissociated PPS-ICBR tumors treated with vehicle, vorinostat, ICB, or ICB+vorinostat. (K) Quantification of CD8α IHC signals for PPS-ICBR tumors in mice treated with vehicle, vorinostat, ICB, ICB+vorinostat, or ICB+vorinostat+anti-CD8. N=7/group. (L) Growth curves of PPS-ICBR tumors in mice treated with vehicle (n=8), ICB+vorinostat+anti-CD8 (n=8), or ICB+vorinostat+isotype (n=5). Arrows indicate the dosing of anti-CD8 or isotype IgG. (M) Individual growth curves of PPS-ICBR tumors (n=6, no growth) or MC38 tumors (n=2, both grew) in C57BL/6 hosts previously cured of PPS-ICBR tumors with ICB+vorinostat treatment. In (D), (K) and (L), data represent mean ± SEM. *P<0.05, **P<0.01, ***P<0.001, ****P<0.0001, ns, not significant; unpaired t-test for (D), Mann-Whitney test for (K), two-way ANOVA with Geisser-Greenhouse correction and Tukey’s multiple comparison test for (L).

Despite the *in vitro* activity, vorinostat monotherapy *in vivo* failed to affect PPS-ICBR tumors (**Fig. 2G**). Because α-tubulin acetyl Lys40 (α-tub-K40Ac) is deacetylated by HDAC6 (a class IIb HDAC inhibited by vorinostat), α-tub-K40Ac is a valid pharmacodynamic marker for vorinostat. Vorinostat indeed augmented α-tub-K40Ac levels *in vitro* and *in vivo*, yet the signals *in vivo* declined rapidly and returned to the baseline 6 hours after dosing (**Fig. 2H**). This result is consistent with the short half-life (about 2-hour oral) of vorinostat in patients ^27^, presumably accounting for its limited efficacy as monotherapy in our model.

Strikingly, vorinostat dramatically enhanced the efficacy of ICB (anti-PD1+anti-CTLA4), with 9 of 24 combo-treated tumors showing complete response (**Fig. 2G**). Vorinostat enhanced H3K9Ac levels in vorinostat-treated and combo-treated tumors (**Fig. 2I**). Moreover, ICB+Vo led to the highest H-2K^b^ level (**Fig. 2J**). At the histological level, whereas the other three groups showed poor CD8^+^ T cell infiltration, the ICB+Vo group was infiltrated with myriad CD8^+^ T cell infiltration (**Fig. 2K**). To verify the essential role of CTLs in the synergistic effect by ICB+Vo treatment, CD8^+^ T cells were depleted with three doses of anti-CD8 antibody when ICB+Vo commenced (**Fig. 2K**), which abrogated the anti-tumor effect of ICB+Vo (**Fig. 2L**). The efficacy from ICB+Vo was also abrogated when *Batf3^null^* mice were used as the hosts. Because *Batf3* deficiency depletes DCs capable of cross-presenting tumor antigens to CD8^+^ T cells ^28^, this result further proves the essential role of CTLs in ICB+Vo efficacy. Finally, we re-challenged 3 mice fully cured by ICB+Vo with PPS-ICBR cells 55 days after the initial start of treatment. Having detected no tumor growth 40 days later, we challenged 1 of the 3 mice with MC38 colorectal cancer cells in both flanks and readily observed engraftment (**Fig. 2M**). In summary, vorinostat upregulated MHC-I expression and various immune-related pathways in PPS-ICBR cells and markedly restored the sensitivity of PPS-ICBR tumors to ICB through the CTL function.

### Vorinostat plus ICB reshapes the tumor immune microenvironment

To examine how vorinostat and ICB altered the TIME, we immunophenotyped the tumors 12 days after the start of the therapy using CyTOF (**Fig. 3A-B**). The CyTOF antibody panel targeted 27 cell surface markers and 10 intracellular markers. Cytobank was used to analyze the CyTOF data where tSNE was used for dimension reduction and cell clusters were annotated based on marker expression patterns (**Fig. 3C**). Both cell cluster percentages and densities showed visually recognizable changes in immune populations associated with the ICB+Vo condition, including the dramatic increase of NK cells and CD8^+^ T cells (purple) and conventional type 1 dendritic cells (cDC1) (dark blue) as well as the disappearance of M2 tumor-associated macrophages (TAMs, cyan) (**Fig. 3D-E**). For T cells, CD8^+^ CTLs and the central memory (CD44^+^ CD62L^+^) and effector memory (CD44^+^ CD62L^-^) subsets increased substantially only in the ICB+Vo group, CD4^+^ helper subset was upregulated in both ICB and ICB+Vo groups, and Treg (CD25^+^ Foxp3^+^) showed no change (**Fig. 3F**). As a result, CD8^+^ effector/Treg ratio peaked in the ICB+Vo condition. For myeloid cells, M1 macrophages (CD11b^+^ F4/80^+^ MHC-II^+^) and M2 macrophages (CD11b^+^ F4/80^+^ MHC-II^-^ CD206^+^) followed the opposite trend ─ ICB+Vo group had the highest M1 and lowest M2 populations, elevating the M1/M2 ratio (**Fig. 3G**). Dendritic cells (DCs, CD11c^+^ MHC-II^+^) also showed highest level in ICB+Vo, whereas Mo-MDSCs (CD11b^+^ Ly6G^-^ Ly6C^high^) or PMN-MDSCs (CD11b^+^ Ly6G^+^ Ly6C^low^) did not change much even by the combination therapy (**Fig. 3G**). PD-1 and PD-L1 expressions were restricted to T cells and myeloid cells, respectively. In summary, ICB+Vo caused most dramatic changes compared with vehicle, whereas ICB caused moderate changes in some immunocytes, and vorinostat had negligible effect.

**Figure 3.**
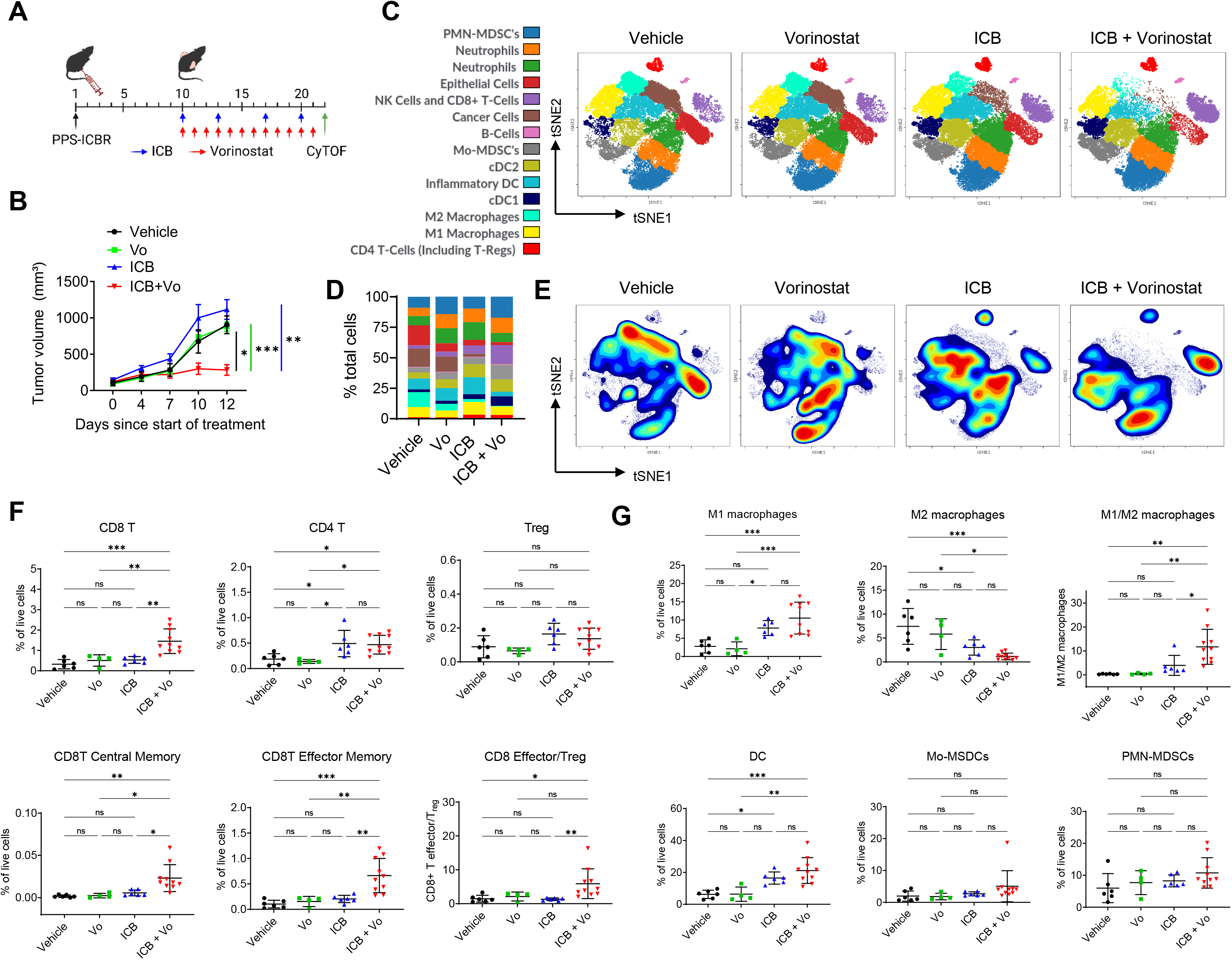
Vorinostat plus ICB reshapes the tumor immune microenvironment. (A) Schematic of the interim cohorts of vorinostat and/or ICB treatments for CyTOF. (B) Growth curves of PPS-ICBR tumors in mice treated with vehicle (n=4), vorinostat (n=4), ICB (n=6), and ICB+vorinostat (n=8). (C) CyTOF tSNE plots showing the 14 cell clusters annotated for the 4 conditions. (D) Cell cluster percentages plotted for the 4 conditions. (E) Cell cluster densities projected on the tSNE maps for the 4 conditions. (F) Percentages of tumor-infiltrating T cell subsets in the 4 conditions. (G) Percentages of tumor-infiltrating myeloid subsets in the 4 conditions. In (B), (F) and (G), data represent mean ± SEM. *P<0.05, **P<0.01, ***P<0.001, ns, not significant; two-way ANOVA with Geisser-Greenhouse correction and Tukey’s multiple comparison test for (B), one-way ANOVA with Tukey’s multiple comparison test for (F) and (G).

To further elucidate the shift in myeloid cells, a second CyTOF experiment with a myeloid-focused antibody panel was conducted for a similar four-group therapeutic experiment. We used tSNE for dimension reduction and Phenograph for cell cluster identification. Among the 20 cell clusters identified by Phenograph, 5 clusters had their fractions in the ICB+Vo therapy over 50% among the four conditions, including CD8^+^ T, BATF^hi^ cDC1, BATF^lo^ cDC1, iNOS^+^ DCs, and an unspecified CD11b^+^ population; by contrast, the 2 clusters with fractions in the ICB+Vo therapy lower than 10% among the four conditions were both M2 macrophages (CX3CR1^high^ and CX3CR1^low^). Among the changes, the dramatic increase of iNOS^+^ DCs (CD11b^+^ CD11c^+^ MHC-II^+^) by the ICB+Vo therapy echoes the previous report on this population being indispensable for effective anti-tumor CD8^+^ T cell therapy ^29^. Overall, the lymphocyte and myeloid landscapes reshaped by the ICB+Vo therapy favor an anti-tumor T cell immunity, explaining the superior tumor-eradiating activity of this combination.

### Cyclic ketogenic diet (CKD) and BHB supplementation sensitize PPS-ICBR tumors to ICB therapy

Based on BHB being an endogenous HDACi^24^, we postulated that KD or BHB could replace vorinostat to synergize with ICB in treating PPS-ICBR tumors. First, the HDACi activity of BHB was confirmed because PPS-ICBR cells treated with 3mM BHB showed higher H3K9Ac and MHC-I, comparable to the effect from 5µM vorinostat (**Fig. 4A-B**). When C57BL/6 mice were fed with standard diet (SD) and KD (**Fig. 4C**), KD generated 10-fold increase in serum BHB compared with SD. The basal BHB level (0.2−0.3mM) and KD-induced BHB level (2−3mM) in mice were consistent with the corresponding values in humans^30, 31^. Continuous KD feeding for 2 weeks delayed tumor growth, yet also led to over 20% weight loss consistent with a recent report ^32^.

**Figure 4.**
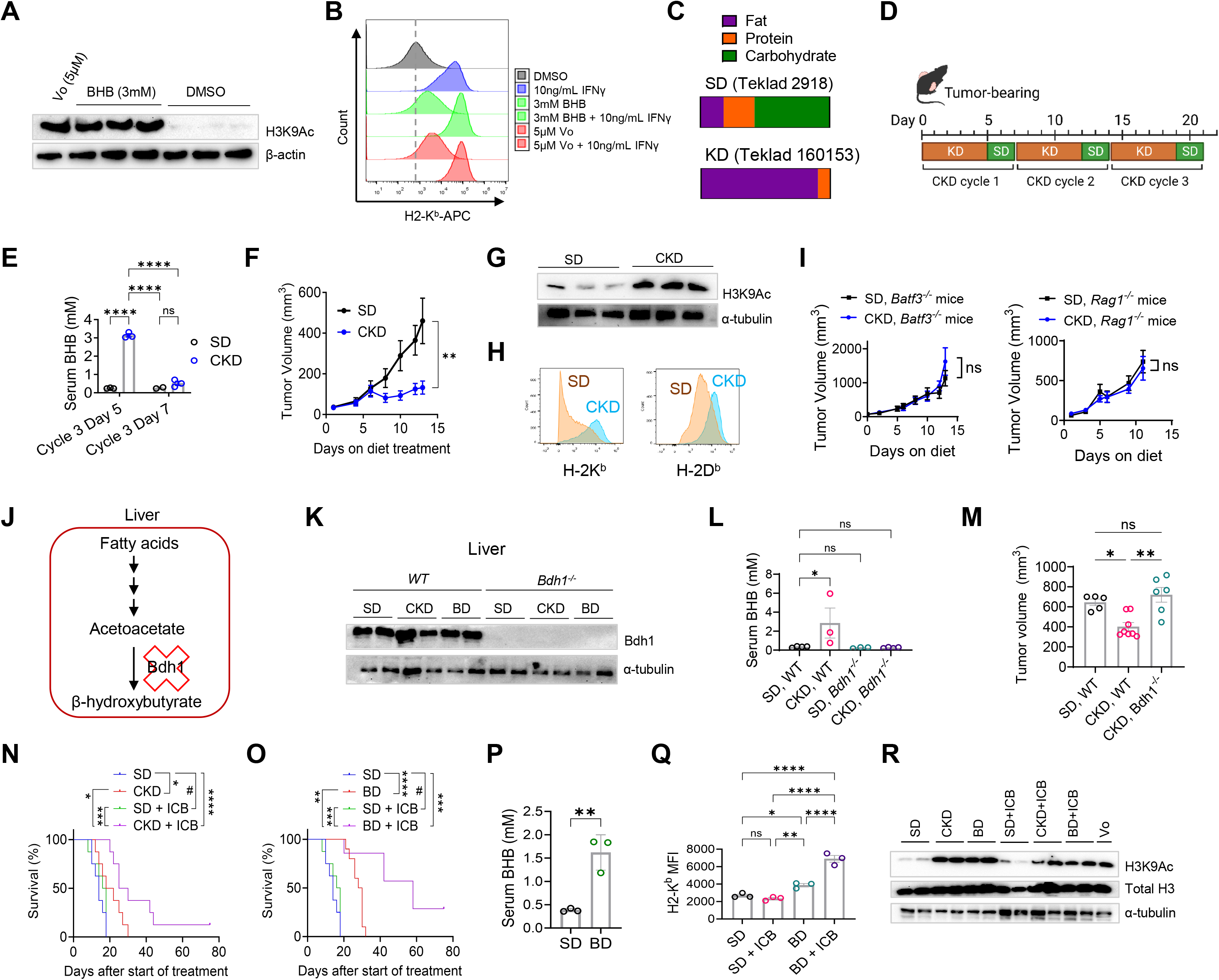
Cyclic ketogenic diet and BHB supplementation sensitize PPS-ICBR tumors to ICB therapy. (A) Western blot of H3K9Ac and β-actin of PPS-ICBR cells treated with 5µM vorinostat, 3mM BHB and DMSO control for 12h. (B) Flow Cytometry of histograms of H2-K^b^ for PPS-ICBR cells treated as indicated for 12h. (C) Caloric composition of SD and KD used in the study. (D) Schedule of CKD feeding. (E) Serum BHB level for C57BL/6 mice fed with SD (n=3) or CKD (n=3) for 3 cycles with BHB measured at cycle 3 day 5 and day 7. (F) Growth curves of PPS-ICBR tumors in C57BL/6 mice fed with SD (n=14) or CKD (n=10). (G) Western blot of H3K9Ac and α-tubulin for PPS-ICBR tumors from mice fed with SD or CKD. (H) Flow cytometry of H2-K^b^ and H2-D^b^ for PCa cells in the dissociated PPS-ICBR tumors from mice fed with SD or CKD. (I) Tumor growth curves of PPS-ICBR subcutaneous tumors in immunocompromised *Batf3^-/-^* or *Rag1^-/-^* C57BL/6 mice fed either a SD or CKD. (*Batf3^-/-^* SD N=8 CKD N=8. *Rag1^-/-^* SD N=6 CKD N=10) (J) The rationale for *Bdh1* knockout to abolish BHB from hepatic ketogenesis. (K) Western blot of Bdh1 and α-tubulin for liver tissue from wild type (WT) or *Bdh1^-/-^* mice fed with SD, CKD, or BD. (L) Serum BHB levels from WT and *Bdh1^-/-^* mice fed with SD or CKD (n=3∼4). (M) Volumes of PPS-ICBR tumors grown in WT mice fed with SD (n=5) or CKD (n=8) or *Bdh1^-/-^* fed with CKD (n=6). (N) Survival curves of PPS-ICBR-bear mice treated as indicated (n=8/group). Tumor volumes at 1000mm^3^ were recorded as the endpoint. (O) Survival curves of PPS-ICBR-bear mice treated as indicated, SD (n=8), BD (n=10), SD+ICB (n=8), or BD+ICB (n=7). Tumor volumes at 1000mm^3^ were recorded as the endpoint. (P) Serum BHB levels in PPS-ICBR-bearing mice fed with SD (n=3) or BD (n=3). (Q) Flow cytometry median fluorescence intensity (MFI) of H2-K^b^ for PCa cells in the dissociated PPS-ICBR tumors from mice treated with SD, BD, SD+ICB or BD+ICB (n=3/group). (R) Western blot of H3K9Ac, total H3 and α-tubulin for PPS-ICBR tumors from mice treated as indicated, with PPS-ICBR cells treated with 5uM vorinostat used as the control lane. In (E) (F) (I) (L) (M) (P) (Q), data represent mean ± SEM. *P<0.05, **P<0.01, ***P<0.001, ****P<0.0001, ns, not significant; two-way ANOVA with Tukey’s multiple comparison test for (E), Mann-Whitney test for (F) (I), one-way ANOVA with Tukey’s multiple comparison test for (L) (M) (Q), log-rank test for (N) (O), unpaired t-test for (P).

To alleviate the adverse effect of continuous KD, we designed a CKD regimen where tumor-bearing mice were fed 5 days on KD and 2 days on SD through a weekly cycling (**Fig. 4D**). For a 7-day cycle, serum BHB reached 3mM at day 5 of the KD period and returned to basal level at day 7 (**Fig. 4E**). CKD not only allowed the mice to gain weight at a comparable pace as those fed with SD, but also delayed tumor growth more effectively than continuous KD (**Fig. 4F**). CKD augmented H3K9Ac signals of PPS-ICBR tumors (**Fig. 4G**), and PCa cells of CKD-fed mice exhibited higher level of H2-K^b^ and H2-D^b^ than PCa cells of SD-fed mice (**Fig. 4H**). The efficacy of CKD was abolished in both *Batf3^-/-^* mice and *Rag1^-/-^*mice (**Fig. 4I**). In brief, CKD had single-agent tumor-decelerating activity, whereas vorinostat did not, and this activity is dependent on antigen cross-presentation by DCs and adaptive immunity of the host.

To elucidate the role of BHB in the anti-tumor effect from CKD, we compared PPS-ICBR tumor growth under CKD in wild-type (WT) mice and mice deficient for 3-hydroxybutyrate dehydrogenase 1 (*Bdh1*) ^33^. Bdh1 is the enzyme responsible for converting acetoacetate (the product from fatty acid oxidation) to BHB during hepatic ketogenesis (**Fig. 4J**). *Bdh1^-/-^* mice were confirmed to lack Bdh1 and serum BHB despite CKD feeding (**Fig. 4K-L**). Remarkably, the effect of CKD on tumor growth was abrogated in *Bdh1^-/-^* mice (**Fig. 4M**), proving the essential role of BHB in ketogenesis-induced tumor retardation.

To determine if ketogenesis enhances ICB, we treated PPS-ICBR-bearing mice with SD, CKD, SD+ICB (anti-PD1+anti-CTLA4), or CKD+ICB. CKD (but not ICB) extended survival as single therapy compared with SD (**Fig. 4N**). CKD+ICB further improved survival (**Fig. 4J**) and reached 14.3% tumor curing. To test the direct effect of BHB on tumor response, we supplemented SD with the ketone diester 1,3-butanediol, a precursor for BHB that is readily converted to BHB *in vivo* through hydrolysis ^34^. Gratifyingly, this 1,3-butanediol-supplemented diet (BD) showed an even more potent effect than CKD to delay tumor growth as a single agent and cured 23.1% of the tumors when combined with ICB (**Fig. 4O**). BD-fed mice reached 1.5mM serum BHB (**Fig. 4P**), a level lower than the 3mM reached by CKD but comparable with the 0.75-1mM level previously reported by BD feeding ^34^. The increased response of BD over CKD may be partly a result of constant BHB presence as opposed to the intermittent presence of BHB by a CKD. H2-K^b^ level on PCa cells was higher in BD-fed mice than in SD-fed mice, and was further enhanced in BD+ICB mice (**Fig. 4Q**). H3K9Ac level was enhanced in tumors treated with all BHB-high conditions, including CKD, BD, CKD+ICB, and BD+ICB (**Fig. 4R**), confirming the HDACi activity of BHB. In summary, CKD or BD raised BHB level, delayed PPS-ICBR tumor growth as a single agent, and sensitized the tumors to ICB therapy with prolonged survival and 10-20% complete response.

### BHB-enhanced immunotherapy bolsters an anti-tumor immune profile

Comparison of the TIME in mice treated with SD, BD, ICB and BD+ICB may reveal mechanisms underlying the synergy between BD and ICB. We dissociated the PPS-ICBR tumors at day 13 of the four conditions (three biological replicates/condition) and profiled viable cells with 10X Genomics scRNA-seq. After quality control, we obtained single-cell transcriptomes from 5749 cells, and biological replicates for each condition were represented unbiasedly. The cells were clustered into 19 clusters encompassing lymphocytes (CD8^+^ T, CD4^+^ T, Treg, NK) and myeloid cells (2 monocyte subsets, 3 DC subsets, 4 macrophage subsets, 3 neutrophil subsets) based on distinct marker expressions (**Fig. 5A-5C**). BD, SD+ICB and BD+ICB each exhibited distinct shifts of cell fractions compared to SD (**Fig. 5D**). BD+ICB showed the highest number of significantly altered cell types including 4 upregulated populations (CD8^+^ T, Vcan^+^ monocytes, M1-macrophages and Nos2^+^ DCs) and 4 downregulated populations (Trem2^+^ M2-macrophages, Hilpda^+^ neutrophils, Cxcr2^+^ neutrophils, mesenchymal cells) (**Fig. 5E**). Among the increased populations, CD8^+^ T cells and M1-macrophages are well-known for their cancer-antagonistic activities. Nos2^+^ DCs (i.e. iNOS^+^ DCs) also expressed *Tnf*, defining them as TNF/iNOS-producing (Tip)-DCs with reported essential role in effectuating anti-tumor CD8^+^ T cell therapy ^29^. Vcan is a marker for tumor-infiltrating monocytes with the potential to differentiate into TAMs or DCs ^35^. On the contrary, all the decreased populations, including M2-macrophages, neutrophil subsets and mesenchymal cells (mainly cancer-associated fibroblasts), often exhibit pro-tumor and immunosuppressive activities in PCa ^36^; hence their decreases reinforce an overall anti-cancer microenvironment. The immune population changes in the BD+ICB condition are reminiscent of the changes observed in the ICB+Vo therapy, such as heightened CD8^+^ T cells, increased M1 and decreased M2 macrophages, and elevated iNOS^+^ DCs. Nonetheless, the conspicuous reduction of neutrophils (Hilpda^+^ and Cxcr2^+^) by BD+ICB and the reduction of Cxcr2^+^ neutrophils by BD alone (**Fig. 5E**) were not observed by either Vo or ICB+Vo, suggesting the existence of HDACi-independent activity of BHB to restrict tumor-associated neutrophils/PMN-MDSCs. This difference should partly explain why BD and CKD delayed tumors and vorinostat did not.

**Figure 5.**
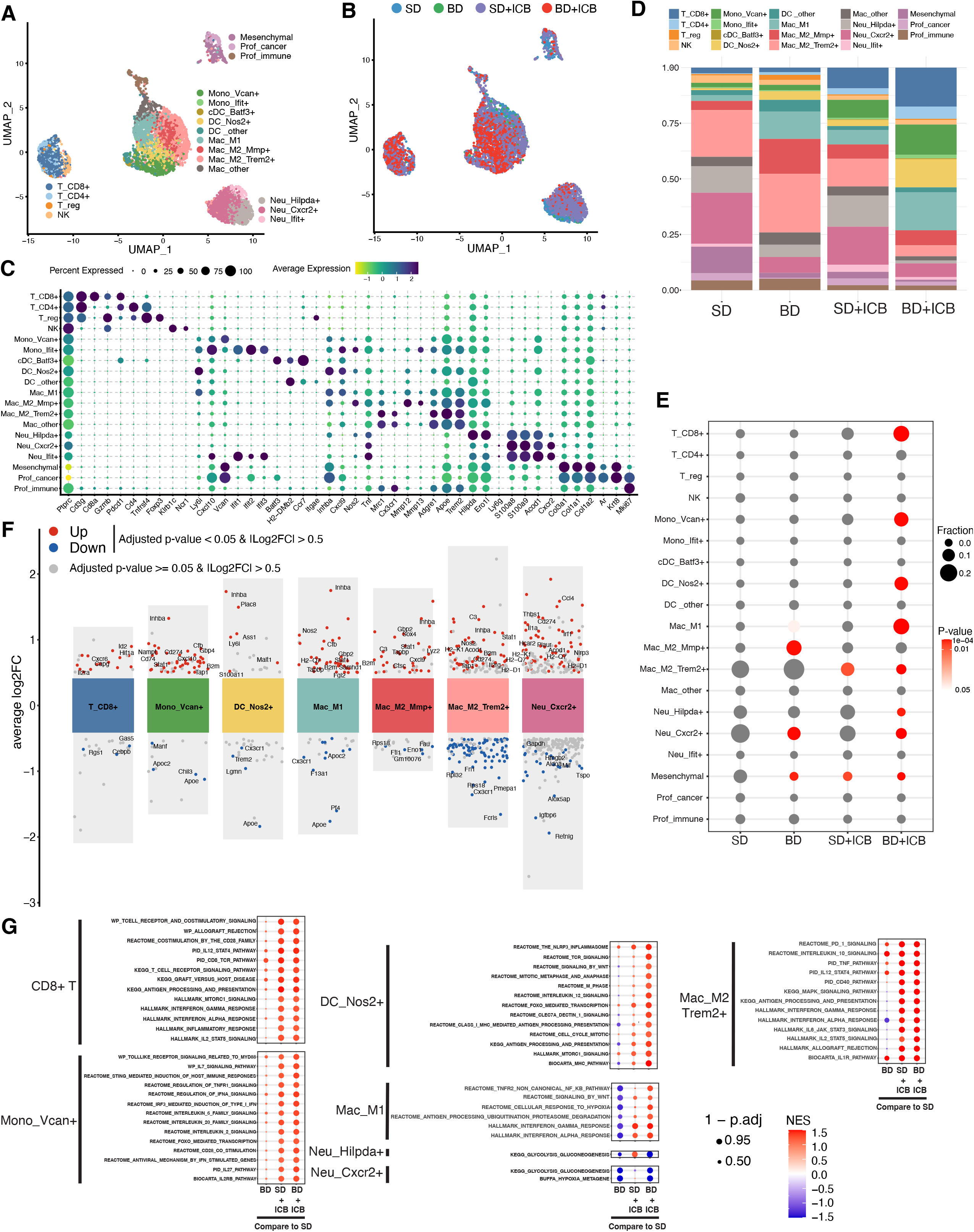
BHB-enhanced immunotherapy bolsters an anti-tumor immune profile. (A) Uniform Manifold Approximation and Projection (UMAP) of 5749 single cells combined from 4 conditions with cells colored by annotated cell clusters. (B) Single cells on the UMAP colored by the four conditions. (C) Dot plot of a selection of marker genes used to annotate the cell clusters. (D) Proportions of the 19 cell clusters for the 4 conditions with colors matching the UMAP. (E) Dot plot of the proportions of the 19 cell clusters for the 4 conditions with fractions denoted with dot sizes and significance levels denoted with a scale of red color. (F) Top differentially expressed genes for CD8^+^ T cells and 6 myeloid subsets between SD and BD+ICB. (G) Dot plot showing the representative pathways significantly enriched for BD+ICB compared with SD for seven cell clusters, with GSEA normalized enrichment scores (NES) denoted by colors (red for positive enrichment, blue for negative enrichment) and adjusted P values denoted by dot sizes. Enrichment results for BD and SD+ICB compared with SD were also plotted.

Besides cell fraction changes, we also examined the differentially expressed genes for the cell clusters between SD and BD+ICB, with the most significant ones plotted for CD8^+^ T cells and 6 myeloid subsets (**Fig. 5F**). CD8^+^ T cells from BD+ICB upregulated several genes essential for cytotoxic functions, including *Id2*, *Hif1a*, *Cxcr6* and *Il2ra* (*Cd25*) ^37–39^. For myeloid cells, the genes simultaneously upregulated in monocytes, macrophages and neutrophils are involved in interferon-γ response and antigen presentation (e.g., *B2m*, *Tap1*, *Irf1*, *Stat1*, *Cxcl10*, *Cd274*), consistent with an expected communication between these myeloid cells with CD8^+^ T cells to induce CTL killing of PCa cells and manifest response of ICB. Several MHC-I genes (*H2-K1*, *H2-D1*, *H2-Q7*) were upregulated in macrophages and neutrophils (**Fig. 5F**), suggesting the stimulation of MHC expression by BHB extends beyond cancer cells. The significant downregulation of Apoe in Vcan^+^ monocytes, M1-macrophages and Nos2^+^ DCs (**Fig. 5F**) was likely releasing immunosuppression and contributing to ICB responsiveness ^40^.

To understand the TIME modulation at the pathway level, we conducted gene set enrichment analysis (GSEA) of MSigDB hallmark and canonical pathways to compare BD, SD+ICB and BD+ICB with SD, respectively (**Fig. 5G**). For CD8^+^ T cells, both SD+ICB and BD+ICB enriched for interferon-α and interferon-γ responses, T cell receptor signaling, IL12-STAT4 pathway, IL2-STAT5 pathway, CD28 co-stimulation pathway, and mTORC1 signaling. For Vcan^+^ monocytes, both SD+ICB and BD+ICB enriched for various interleukin pathways (IL2, IL6, IL7, IL20, IL27), type I interferon (interferon-α)/IRF3 signaling, Toll-like receptor signaling, STING pathway, and CD28 co-stimulation pathway. Hence, SD+ICB and BD+ICB shared marked similarities in pathway activations for CD8^+^ T cells and Vcan^+^ monocytes whereas BD alone had little effect, suggesting ICB’s central role in stimulating these pathways. Therefore, the significance of BD in the BD+ICB combination is more likely through mitigating immunosuppression (e.g., reducing Cxcr2^+^ neutrophils). The comparable effect of SD+ICB and BD+ICB on pathway enrichment was also observed for Trem2^+^ M2-macrophages (**Fig. 5G**), consistent with these cells being reduced in proportion by both treatments (**Fig. 5E**). Nos2^+^ DCs (Tip-DCs) under BD+ICB enriched for pathways relevant to proliferation (mitotic metaphase and anaphase, M phase, cell cycle mitotic), activation (IL12, mTORC1, Wnt, FOXO, NLRP3 inflammasome) and antigen-presenting (class I MHC, antigen processing and presentation, TCR signaling) (**Fig. 5G**), suggesting the active participation of these cells in presenting tumor antigens and bridging T cell immunity. Interestingly, simultaneous BD and ICB treatments were required for elevating and activating Nos2^+^ DCs as neither BD nor SD+ICB could raise the fraction or enrich pathways for Nos2^+^ DCs. M1-macrophages also relied on the BD+ICB combination to activate pro-inflammatory pathways, such as for interferon-α and interferon-γ responses and NFκB pathway, and the combination appeared capable of reversing the negative influence of BD alone on these pathways (**Fig. 5G**). Finally, BD and BD+ICB weakened glycolysis and gluconeogenesis pathways in Hilpda^+^ and Cxcr2^+^ neutrophils (**Fig. 5G**), suggesting BHB-induced metabolic debilitation of specialized neutrophils. Overall, scRNA-seq reveals that while BD and ICB each generated a moderate impact on the immune landscape, the combination drastically reprogrammed the TIME and fostered a CD8^+^ T-dominated adaptive immune response to eliminate cancer cells.

### BD+ICB therapy augments Cxcr3/Cxcr6 signaling for T cells and strengthens myeloid differentiation

To explore how BHB and ICB therapies modulate intercellular communications, we used CellChat to quantitatively infer cell-cell communication networks based on ligand-receptor expression patterns from scRNA-seq data ^41^. First, interactomes for each condition were depicted with circle plots for CD8^+^ T, CD4^+^ T (Treg excluded), monocytes (Vcan^+^), DCs (Nos2^+^), macrophages (M1, Trem2^+^ M2, Mmp2^+^ M2), neutrophils (Hilpda^+^, Cxcr2^+^), and mesenchymal cells (**Fig. 6A**). SD+ICB and BD+ICB conditions featured visually more extensive connections among cell clusters, especially feeding in and out of CD8^+^ and CD4^+^ T cells (including autocrine loops), consistent with the anticipated activity of ICB to reinvigorate effector T cells. When displayed on scatter plots to compare the outgoing and incoming interaction strengths for all the 19 cell clusters across the conditions, SD+ICB and BD+ICB also showed dramatically augmented counts of sent or received signals for CD8^+^ T, CD4^+^ T, monocytes (Vcan^+^, Ifit^+^), macrophages (M1, Trem2^+^ M2, Mmp^+^ M2) and DCs (Nos2^+^), yet Treg and Batf3^+^ cDCs remained weak in the communications. The information flow of each signal molecule charted in CellChat also demonstrated the augmented intercellular communications under SD+ICB and BD+ICB conditions, with BD+ICB selectively enriched for a few signals, including IL2, IL4, VEGF, TRAIL, LIGHT, and ACTIVIN.

**Figure 6.**
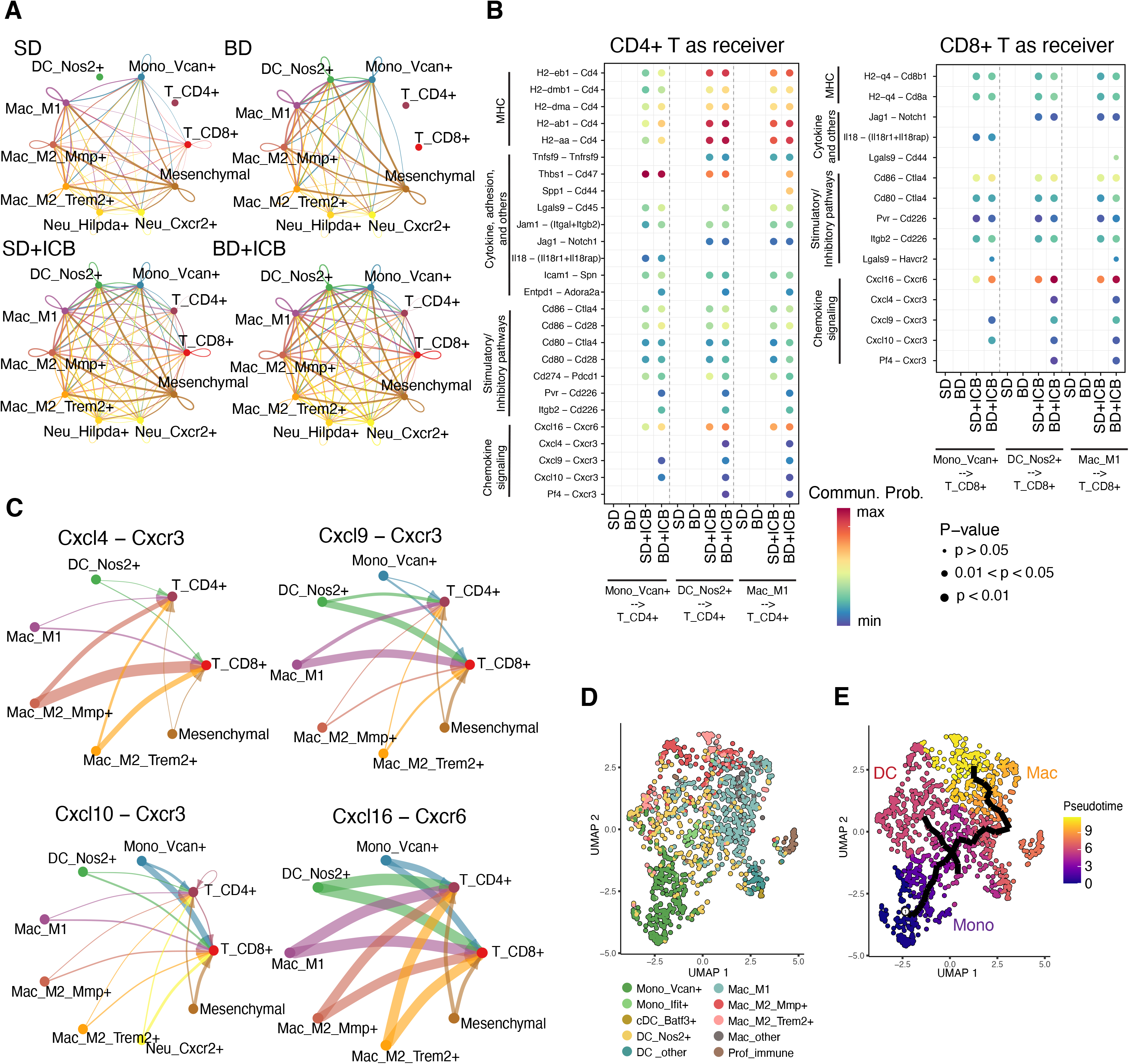
BD+ICB therapy augments Cxcr3/Cxcr6 signaling for T cells and strengthens myeloid differentiation. (A) Circle plots showing the strengths of intercellular interactions in each treatment condition for 10 cell clusters. (B) Dot plot showing LRPs with significant communication probabilities between at least one of the three myeloid populations and CD4^+^ or CD8^+^ T cells under BD+ICB condition. Dot colors denote communication probability scales and dot sizes denote P values (P<0.05 for all dots shown). Corresponding dots (or the lack thereof) for SD, BD and SD+ICB conditions were also plotted. (C) Circle plots showing the strengths of four LRPs among defined myeloid populations, mesenchymal cells, CD8^+^ and CD4^+^ T cells, with arrows pointing to the signal receivers. (D) Myeloid-focused UMAP (excluding neutrophils) created by Monocle 3, with cell cluster annotations and colors carried from the original UMAP in Figure 5A. (E) Pseudotime trajectory inferred by Monocle 3 overlaid on the myeloid-focused UMAP. The origin position 1 was set near Vcan^+^ monocytes. Branch ends were labeled based on the locally enriched myeloid lineages: monocytes (Mono), DC, and macrophages (Mac).

To gain insight into how myeloid cells regulate T cells, we focused on the ligand-receptor pairs (LRPs) with myeloid cells (Vcan^+^ monocytes, Nos2^+^ DCs, M1-macrophages) as the ligand producers (“sender”) and CD8^+^ or CD4^+^ T cells as the receptor owners (“receivers”). The significant LRPs for the BD+ICB condition were clustered based on functional categories, including MHC-related, T cell stimulatory/inhibitory signaling, chemokine signaling, and others (e.g., cytokines, adhesion) (**Fig. 6B**). A large proportion of these LRPs were also significantly represented in the SD+ICB condition, albeit the communication probabilities between the two conditions can vary (e.g., Cxcl16-Cxcr6 was stronger in BD+ICB than SD+ICB for all three myeloid populations toward both CD8^+^ and CD4^+^ T cells, Cd274-Pdcd1 was weaker in BD+ICB than SD+ICB for all three myeloid populations toward CD4^+^ T cells). Some LRPs were only present in BD+ICB, suggesting their specific contribution to the tumor-killing T cell activity by BD+ICB therapy, with the most outstanding example being the several LRPs with Cxcr3 as the receptor (Cxcl4-Cxcr3, Cxcl9-Cxcr3, Cxcl10-Cxcr3) (**Fig. 6B**). Circle plots for the Cxcr3- and Cxcr6-oriented LRPs under BD+ICB condition depicted the strong and directional connections from myeloid and mesenchymal cells toward CD8^+^ and CD4^+^ T cells (**Fig. 6C**), consistent with the critical role of Cxcr3 and Cxcr6 to recruit and sustain CTLs to control tumors ^42, 43^. Interestingly, while the Cxcr3 chemokine Cxcl9 originated mainly from Nos2^+^ DCs and M1-macrophages, another Cxcr3 chemokine Cxcl10 predominantly emerged from Vcan^+^ monocytes (**Fig. 6C**), highlighting the cell type-specific chemokine production. The lack of LRP enrichment in SD and BD conditions (no dots in Fig. 6B) indicates the absence of meaningful crosstalk to T cells when immune checkpoint pathways were unperturbed.

The simultaneous increase of Vcan^+^ monocytes, Nos2^+^ DCs and M1-macrophages by BD+ICB suggests relationships among these cells. We focused on the myeloid island in the middle of the UMAP and applied an unsupervised inference method, Monocle 3 ^43^, to construct the potential transitional trajectories among myeloid cells under BD+ICB condition (**Fig. 6D**). Pseudotime suggested a trajectory from monocytes to both DC and macrophage branches (**Fig. 6E**), consistent with the ability of monocytes to differentiate into TAMs and monocyte-derived DCs (moDCs) in the TIME ^44^. In summary, BD+ICB treatment is associated with augmented cell-cell communications underscored by Cxcr3 and Cxcr6-mediated T cell engaging and enhanced myeloid differentiation from monocytes.

## Discussion

Our study connects two promising anti-cancer regimens, HDACi and KD, demonstrating that HDACi plus ICB and KD plus ICB generate synergistic anti-tumor efficacy in a newly developed ICB-refractory PCa model. Therefore, this study features significant translational implications for treating advanced PCa with ICB, a notoriously tricky clinical challenge. Regarding combining HDACi and ICB, multiple early phase clinical trials are testing the combination of vorinostat and anti-PD1 in malignancies such as glioblastoma (NCT03426891), urothelial and renal carcinomas (NCT02619253), non-small cell lung cancer (NCT02638090), and lymphomas (NCT03150329). Early reports of several trials showed that pembrolizumab plus vorinostat was well tolerated with promising anti-tumor activities ^45–47^. Vorinostat used in our study (25 mg/kg daily in mice) is equivalent to 2 mg/kg for human patients with a reference body weight 60 kg based on the mouse-human conversion factor 12.3 (FDA guidance ^48^). This dose is significantly lower than the recommended vorinostat dose to treat adult cutaneous T cell lymphoma (400 mg/day) and falls below or within the lower spectrum for the clinical trials that combine vorinostat and anti-PD1 (e.g., NCT02638090 used vorinostat at 200 or 400 mg/day to combine with pembrolizumab ^45^). Therefore, the vorinostat dose based on our preclinical results should be tolerated if translated to clinical trials.

Regarding using KD as cancer therapy, KD as monotherapy and as an adjuvant for chemotherapy or hormonotherapy has been investigated in clinical trials for various tumor types, including rare cancers ^49^. Unfortunately, many trials were plagued by a lack of patient compliance due to poor palatability of KD and sometimes side effects. Therefore, the clinical translation of the CKD and BD (BHB supplement) from our study would represent attractive solutions to addressing compliance issues. Compared with previous preclinical studies that used oral gavage or intraperitoneal administration of BHB or BHB esters ^50, 51^, 1,3-butanediol supplementation into a regular diet should further facilitate compliance. One clinical trial was initiated recently to test the combination of KD (continuous or discontinuous) or BHB supplementation with nivolumab plus ipilimumab to treat metastatic renal cell carcinoma (NCT05119010). The result from this trial is eagerly awaited to provide evidence for ketogenesis-enhanced immunotherapy.

Two recent studies reported the effect of KD to enhance ICB therapy in preclinical models ^50, 52^. While our study agrees with the overall conclusion of these two publications, a few unique strengths exist in our study. (1) We are the first to demonstrate ketogenesis-enhanced immunotherapy in PCa models. Dai and Ferrere demonstrated the enhancement of PD1 or CTLA4 antibodies (but not both) by KD in syngeneic models of melanoma, renal cancer and colon cancer ^50, 52^. Compared to these cancer types, PCa exhibits unique challenges in ICB resistance, accentuated by its low response to dual anti-PD1 and anti-CTLA4 inhibition ^53^. By modeling dual ICB therapy resistance, our results based on the PPS-ICBR model address a critically unmet need. (2) scRNA-seq was applied in our study to explore the TIME reprogramed by BD and ICB, which revealed the dramatic upregulation of CD8^+^ T cells and three myeloid populations (Vcan^+^ monocytes, Nos2^+^ DCs, M1-macrophages) only under the BD+ICB treatment. (3) Regarding the mechanisms, while Dai et al. discovered that KD-triggered energy deprivation stimulated AMPK activation which further caused PD-L1 degradation and interferon upregulation ^52^, Ferrere et al. associated the KD-enhanced immunotherapy to compositional changes of gut microbiota ^50^. By contrast, our result emphasizes both cancer-cell-intrinsic and extrinsic mechanisms elicited by BD+ICB. At the cancer-cell-intrinsic level, the pan-HDACi activity of BHB augmented MHC-I expression in PCa cells, rendering them more visible to the immune system. At the cancer-cell-extrinsic level, similar to vorinostat, BD alone had a relatively moderate effect on the TIME composition (except for reducing Cxcr2^+^ neutrophils). However, the BD+ICB combination drastically reshaped the TIME to favor tumor-killing T cell immunity, highlighted by heightened CD8^+^ T cells and shifted myeloid infiltration and differentiation (monocytes toward antigen presenting cells, elevated M1/M2 macrophage ratios, and diminished neutrophils). While most immunocyte changes were rightfully consistent between BD+ICB and Vo+ICB combinations, a notable difference was that BD+ICB depleted Hilpda^+^ neutrophils and Cxcr2^+^ neutrophils, but Vo+ICB did not. GSEA suggests that these neutrophils may be compromised due to the diminished glycolysis and gluconeogenesis pathway, a metabolic vulnerability hit by the BD+ICB but not the Vo+ICB condition. We believe these intrinsic and extrinsic mechanisms at least complement the mechanisms in Dai and Ferrere’s studies and illustrate how an optimized KD regimen can take diverse routes to enable effective immunotherapy.

Our experiments using *Bdh1^-/-^*, *Batf3^-/-^*and *Rag1^-/-^* mice demonstrated the essential role of both BHB and adaptive immunity in establishing the anti-tumor activity of CKD. This result echoes recent studies that elucidated various immune-modulatory activities of BHB in various physiological contexts, which can be classified as T cell-related and myeloid cell-related. For T cells, KD was shown to expand γδ T cells in the mouse lung and improve antiviral resistance against influenza virus ^54^. BHB reduced the mortality of mice infected with SARS-CoV-2 by fueling oxidative phosphorylation and restoring the function of CD4^+^ T cells ^55^. BHB enhanced human T cell responses and memory T cell formation *in vitro* ^56^. Therefore, KD/BHB generally upholds T cell functions, which was consistent with the enhanced Cxcr3/Cxcr6 signaling and increased CD8^+^ T cell infiltration by BD+ICB treatment in our model, although this increase depended on the presence of both BD and ICB. For myeloid cells, BHB often plays an anti-inflammatory role. BHB blocked activation of the NLRP3 inflammasome by inhibiting potassium efflux in monocytes/macrophages ^57^, and abrogated NLRP3/caspase 1-dependent IL1β secretion from neutrophils to alleviate gout flare ^58^. In an alcoholic hepatitis model, BHB supplementation signaled through its receptor Hcar2 (hydroxycarboxylic acid receptor 2) to increase M2-macrophages and reduce neutrophil influx leading to less liver damage ^59^. In a glioblastoma model, KD increased M2 and decreased M1-macrophages, justifying the combination of KD and CSF1R inhibitor ^60^. Interestingly, we also observed that BD alone increased Mmp^+^ M2-macrophages and decreased Cxcr2^+^ neutrophils (Fig. 5E). Intriguingly, the addition of ICB curbed BD’s effect on Mmp^+^ M2-macrophages yet amplified BD’s effect on Cxcr2^+^ neutrophils (Fig. 5E), with the net result supporting anti-tumor immunity. Future work will further resolve why the interplay between BD and ICB generates an immune landscape maximizing CTL activity and minimizing immunosuppression.

In summary, we developed a syngeneic PCa cell line series with PPS-ICBR being a robust model for dual ICB resistance. Using this model, we demonstrated that epigenetic modulation with pharmacological HDACi vorinostat or endogenous HDACi BHB (via CKD or BD) sensitized ICB-refractory PCa to PD1/CTLA4 dual ICB therapy. The two combination therapies share underlying mechanisms such as cancer cell MHC-I upregulation and concerted innate and adaptive immune landscape remodeling, culminating in unleashed CTL reinvigoration and tumor eradication. Among the strategies, 1,3-butanediol-supplemented-diet may represent the most attractive option to enhance ICB therapy in patients due to its high chance of patient compliance, low cost, and minimal toxicity.

## Acknowledgments

We thank the Lu lab members for their essential comments and suggestions during this work. We thank Daniel P. Kelly from University of Pennsylvania for sharing Bdh1^-/-^ mice. We appreciate the technical assistance on scRNA-seq from Qingfei Wang and Siyuan Zhang. We are grateful for the support from core facilities used in this study, especially Freimann Life Science Center (Teri Highbaugh) and Genomics and Bioinformatics Core Facility (Michael Pfrender, Melissa Stephens, Jacqueline Lopez, Brent Harker). This work was supported by a research grant from American Institute for Cancer Research (Xin Lu), National Institutes of Health grant 5F99CA274694 (Sean Murphy), and a core facility grant from Indiana Clinical and Translational Sciences Institute and National Institutes of Health grant UL1TR002529 (Xin Lu). Other support included National Institutes of Health grants R01CA248033 and R01CA280097 (Xin Lu), Department of Defense grants W81XWH2010312, W81XWH2010332, and HT94252310010 (Xin Lu), and Boler Family Foundation (Xin Lu) at University of Notre Dame.

## Author Contributions

S. Murphy: Conceptualization, investigation, methodology, data curation, formal analysis, validation, visualization, funding acquisition, manuscript writing. S. Rahmy, D. Gan: Investigation, methodology, data curation, software, formal analysis, validation, visualization, manuscript writing. Y. Zhu, M. Manyak: Investigation and methodology. J. Li, Xuemin Lu: Supervision. Xin Lu: Conceptualization, investigation, formal analysis, resources, project administration, supervision, funding acquisition, manuscript writing.

## Methods

### Animals

All animal work performed in this study was approved (19-09-5563 and 22-08-7344) by the Institutional Animal Care and Use Committee (IACUC) at University of Notre Dame. All animals were maintained under pathogen-free conditions and cared for per the International Association for Assessment and Accreditation of Laboratory Animal Care policies and certification. C57BL/6J (RRID:IMSR_JAX:000664), Batf3^-/-^ (RRID:IMSR_JAX:013755), Rag1^-/-^(RRID:IMSR_JAX:002216) mice were purchased from Jackson Laboratory and bred in house. Bdh1^-/-^ allele was a generous gift from Daniel P. Kelly at University of Pennsylvania. Only male mice were used for experiments, given the gender association of prostate cancer. All the mice were in the C57BL/6 background.

### Cell lines

PPS-6239, PPS-PD1R, PPS-CTLA4, PPS-ICBR, and MC38 were cultured in DMEM (GE Healthcare, SH30243.FS) supplemented with 10% fetal bovine serum (FBS; GE Healthcare, SH30396.03) and 100U/ml penicillin-streptomycin (Cytiva, SV30010). RM9 was purchased from ATCC (CRL-3312, RRID: CVCL_B461) and cultured in DMEM/F12 (VWR, 45000-344) supplemented with 10% FBS and 100U/ml penicillin-streptomycin. All the cell lines were cultured at 37°C in a humidified incubator with 5% CO2. All cells were tested for mycoplasma-free status using a Mycoplasma Assay Kit (Agilent Technologies, 302109).

### Animal experiments for therapeutic and dietary interventions

Syngeneic tumors were generated by injecting 5×10^5^ tumor cells subcutaneously into both flanks of 6-8 weeks old male mice. When tumors reached 50-100mm^3^, mice were randomized to receive therapeutics, including anti-PD1 (BioLegend, 114116) and anti-CTLA4 (BioLegend, 106207) at 10mg/kg each i.p. twice a week, or vorinostat (MedChemExpress, HY-10221) at 25 mg/kg i.p. daily. For CD8^+^ T cell depletion, tumor-bearing mice received i.p. injection of three doses of 200μg anti-CD8 (BioXCell, BE0061) at the indicated time points. For dietary interventions, the randomized mice were fed ad libitum with standard chow (Teklad, 2918), or ketogenic diet (Teklad, 160153) replenished daily, or cyclic ketogenic diet which followed a weekly cycles with five days on daily ketogenic diet (Teklad, 160153) then two days on standard chow (Teklad, 2918), or 1,3-butanediol-supplemented diet which was prepared by mixing 80ml of 1,3-butanediol (Sigma-Aldrich, 309443), 120ml of water, 198g of standard chow (Teklad, 2918) and 2g of Saccharine (Sigma-Aldrich, 109185), replenished every other day. Mice on the KD were given extra water due to the metabolic requirements of fatty acid degradation. All treatments were continued until the specified experimental endpoints were reached.

### Cell treatment in vitro

PPS-ICBR cells grown in the regular medium were treated with 5µM vorinostat (MedChemExpress, HY-10221), 1:2000 diluted DMSO (Sigma-Aldrich, D4540), or 10ng/ml mouse IFN-γ (Sigma-Aldrich, I4777). For β-hydroxybutyrate treatment, the medium was removed once cells in the regular medium reached ∼70% confluence. Cells were washed once with warm PBS and changed to a medium of DMEM without glucose (VWR, 97060-876), 10% FBS, and 1% Pen/Strep. Glucose was restored to 5mM to mimic the human blood glucose level by diluting from a 500mM stock of glucose (Sigma-Aldrich, G8270). β-Hydroxybutyrate (Cayman Chemical, 14148) was added to 3mM, and the medium was titrated to pH 7.2 using HEPES (VWR, 16777-032). The treatments were maintained for indicated durations before cells were harvested for subsequent experiments.

### Western Blot

Cells and fresh tissue samples were lysed in Radioimmunoprecipitation assay buffer (RIPA) supplemented with protease inhibitors (Bimake, B14012) and phosphatase inhibitors (Bimake, B15002). The western blot procedure has been described previously ^61^. The primary antibodies were used to detect β-actin (Santa Cruz, sc-47778), α-tubulin (Santa Cruz, sc-5286), acetyl-Histone H3 Lys9 (Cell Signaling Technology, 9649), Histone H3 (Cell Signaling Technology, 4499), acetyl-α-tubulin Lys40 (Cell Signaling Technology, 5335), BDH1 (ProteinTech, 15417-1-AP). The secondary antibodies include HRP-conjugated goat anti-mouse (Cell Signaling Technology, 7076) and HRP-conjugated goat anti-rabbit (Cell Signaling Technology, 7074). Signals were detected with Clarity Max ECL Substrate (Bio-Rad, 1705062).

### Immunohistochemistry

Animal tissues were fixed overnight in 10% formalin and embedded in paraffin. Antigen retrieval was performed by heating in a pressure cooker at 95°C for 30 min, followed by 115°C for 1 min in citrate-unmasking buffer (pH 6.0). IHC staining with CD8α antibody (Cell Signaling Technology, 98941) was performed using the VECTASTAIN Elite ABC-HRP Kit (Vector Laboratories, PK-6101) with signals detected with DAB Substrate Kit (Vector Laboratories, SK-4100). Counterstain was performed using hematoxylin (VWR, 10143-608). The IHC slides were scanned using an Aperio ScanScope (Leica). Signals were quantified using functions in ImageJ Fiji.

### Serum β-hydroxybutyrate quantification

Mouse blood (100ul) was collected with submandibular bleeding. Serum β-hydroxybutyrate concentration was measured following the manual instructions of the β-hydroxybutyrate colorimetric detector kit (Cayman Chemical, 700194).

### Bulk RNA profiling and data analysis

Cancer cells treated with DMSO or vorinostat for 12 hours were used to extract total RNA using RNeasy Kit (Qiagen). RNA samples were profiled on the Mouse Genome 430 2.0 Array (Affymetrix) at the Genomics Core Facility at the MD Anderson Cancer Center. The data were analyzed using Transcriptome Analysis Console (TAC) software (Thermo Fisher) to generate a list of differentially expressed genes with cutoff FDR < 0.05, absolute fold change ≥3. Pathway enrichment was performed using Enrichr (RRID: SCR_001575).

### Flow cytometry

Flow cytometry samples were prepared as described previously ^61^ and run on CytoFLEX (Beckman Coulter). Data were analyzed using FlowJo v10.8 (FlowJo, RRID: SCR_008520). Fluorochrome-conjugated antibodies included H2-K^b^-APC (BioLegend, 116518), H2-D^b^-PE/Cyanine7 (BioLegend, 111516).

### CyTOF and data analysis

Tumors were minced into homogenate and rotated at 37°C in dissociation media, DMEM with 10% FBS and 1 mg/ml collagenase IV (STEMCELL Technologies, 07427) for 1 h, followed by passing through 40μm strainers. Erythrocytes were depleted via hypotonic lysis. The CyTOF procedure has been described previously ^61^. The samples were run with Helios CyTOF mass cytometer (Fluidigm) in the Flow Cytometry and Cellular Imaging Core Facility at the MD Anderson Cancer Center. For the first CyTOF experiment (Figure 3), data analysis was performed with Cytobank (Beckman Coulter). Samples were concatenated based on treatment types and then uploaded to Cytobank. tSNE plots of single vialbe CD45^+^ cells were generated using optSNE with 5000 iterations and a KL divergence of 4.329796. FlowSOM was used with a pre-determined number to create 14 cell clusters manually annotated based on marker expression patterns. For the second myeloid-focused CyTOF experiment, data analysis was performed with FlowJo v10.8 (FlowJo, RRID: SCR_008520). After gating cell populations manually, CD45^+^ cells were used for dimension reduction using tSNE with iterations = 1,000, perplexity = 30, and learning rate = 2,800. The Approximate (random projection forest - ANNOY) KNN algorithm was used with the FFT interpolation gradient algorithm. Cell clustering was performed using the FlowJo plugin tool PhenoGraph with parameters K = 30 and Run ID = auto, with clusters manually annotated based on marker expression patterns. We used the plugin tool ClusterExplorer to generate the intensity heat map table of proteins for each condition.

### Single-cell RNA-seq and data analysis

Tumors were minced and dissociated with a mouse dissociation kit (Miltenyi Biote,130-096-730) at 37℃ for 1 hour. Dissociated tumor single-cell suspension was passed through 40μm strainers followed by red blood cell lysis. Then the dead cell was removed using EasySep Dead Cell Removal (Annexin V) Kit (STEMCELL Technologies, 17899) followed by multiplexing with hashtag labeling of TotalSeq-A0303 (BioLegend, 155805), TotalSeq-A0304 (BioLegend, 155807), TotalSeq-A0305 (BioLegend, 155809), TotalSeq-A0306 (BioLegend, 155811), TotalSeq-A0307 (BioLegend, 155813), and TotalSeq-A0308 (BioLegend, 155815). At this step, the samples were transferred to Genomics and Bioinformatics Core Facility at University of Notre Dame, where cells were counted, evenly combined, and loaded onto the Chromium Controller (10X Genomics). The samples were then processed following the manuals of the Chromium Single Cell 3’ Reagent Kits. Next, the samples were sequenced with Illumina NovaSeq 6000 platform at the Medical Genomics Center at Indiana University.

Barcode processing, alignment, filtering, and UMI counting were performed using the Cell Ranger analysis pipeline (v. 6.1). Sample demultiplexing was performed using the CITE-seq-Count pipeline (v. 1.4). Cells with larger than 500 UMI counts in the demultiplexing step were used for downstream analysis in R (v. 4.1). Data pre-processing, normalization, and clustering was performed using the R package Seurat (v. 4.1). Single-cell transcriptomes were initially filtered for quality control. Cells with fewer than 200 genes detected and genes expressed in fewer than three cells were removed. We also filtered cells with the percent of mitochondrial counts larger than 5%. The resulting expression matrix contained 57,49 cells by 18,848 genes, with 1,215 cells from the SD treatment, 490 cells from the BD treatment, 2314 cells from the SD+ICB treatment, and 1730 cells from the BD+ICB treatment. Data normalization was performed using the function NormalizeData with normalization.method = “LogNormalize” and scale.factor = 10000. 2,000 variable genes were chosen using the function FindVariableFeatures with selection.method = “vst.” Data scaling was performed using the ScaleData function. Dimensional reduction was accomplished by performing principal component analysis (PCA) and then using the first 50 principal components for Uniform Manifold Approximation and Projection using default parameters associated with the RunUMAP function. Unsupervised clustering was finished by constructing a shared nearest neighbor (SNN) graph using the FindNeighbors function and then performing graph-based clustering using the “Louvain” algorithm with resolution = 1.8, resulting in 22 clusters. We merged clusters based on their marker genes, leading to 19 cell types in our analysis. Differential expression analysis between clusters was performed using a Wilcoxon rank sum test by the FindAllMarkers function. The dot plots and feature plots for chosen marker genes were obtained by the DotPlot and FeaturePlot functions, respectively. The expression heatmap was generated using the function DoHeatmap.

Signaling pathway gene signatures were curated from the hallmark gene sets, C2, C6, C7, and C8 collections within MSigDB. Normalized enrichment scores (NES) were calculated using the fgsea package v. 1.20, with parameters minSize = 15 and maxSize = 500. Cell-cell communication analysis for each treatment group was conducted using the CellChat package v. 1.4. The analysis followed the standard pipelines provided by the CellChat package, using the ligand-receptor database curated for mouse tissues. We computed the communication probability for each treatment condition using the computeCommunProb function and filtered out the cell-cell communications if there are less than ten cells in certain cell groups, using the filterCommunication function with parameter min.cells = 10. We then inferred the cellular communication network and aggregated cell-cell communication, using computeCommunProbPathway and aggregateNet functions, respectively. The netVisual_circle function generated the circle plots for each condition. The bubble plots with multiple functions were created with the netVisual_bubble function. The circle plots for individual ligand-receptor pairs were created with the netVisual_individual function. The violin plot for chosen genes under different cell types and conditions was generated by the plotGeneExpression function. Visualizations of the dominant senders and receivers for all the conditions were generated with the netAnalysis signalingRole scatter function. The comparison of the overall information flow of signaling pathways among all the conditions was made by the rankNet function.

The pseudotime analysis was conducted using the monocle3 R package v. 1.3.1. We followed the recommended pipelines from the monocle tutorials. Ten cell types from the BD+ICB condition were analyzed. The dimension reduction was performed using the preprocess cds and reduce dimension functions. The trajectory was learned using the learn graph function with parameter close_loop = FALSE. We further inferred the pseudotime using the order cells function and chose the monocyte cell population as the root point.

### Statistical Analysis

*In vitro* and *in vivo* experiments were performed three or more times and conclusions were drawn only when the results were reproducible. One representative result among the replicates was shown in the figures. For non-omics data, statistical analyses were performed using GraphPad Prism v9.3 (RRID: SCR_002798). All data are presented as mean ± SEM (standard error of the mean). We followed this workflow for statistical testing: Shapiro-Wilk test was performed to assess for normality of data distribution: (i) in case of normality, when only two conditions were to test, we performed unpaired t-test; when more than two conditions were to compare, we performed a parametric one-way or two-way ANOVA followed by post hoc test with recommended correction for multiple comparisons to assess the significance among pairs of conditions. (ii) in case of non-normality, when only two conditions were to test, we performed a Mann-Whitney U test; when more than two conditions were to compare, we performed a non-parametric one-way ANOVA followed by a recommended test to assess the significance among pairs of conditions. For survival data, log-rank test was used. Sample sizes, error bars, P values, and statistical methods are noted in the figure legends. Statistical significance was defined as P < 0.05.

### Data and materials availability

Microarray data of DMSO or vorinostat-treated PPS-ICBR cells are available in the Gene Expression Omnibus (GEO) (RRID: SCR_005012) with accession number GSE205469. scRNA-seq data are available at GEO with accession number GSE206561. All the materials generated in this study are available from the corresponding author upon reasonable request.

